# Adaptive strategies under prolonged starvation and role of slow growth in bacterial fitness

**DOI:** 10.1101/2021.12.14.472581

**Authors:** Pabitra Nandy

## Abstract

Adaptive evolution has the power to illuminate genetic mechanisms under a pre-defined set of selection factors in a controlled environment. Laboratory evolution of bacteria under long-term starvation has gained importance in recent years because of its ability to uncover adaptive strategies to overcome prolonged nutrient limitation- a condition thought to be encountered often by natural microbial isolates. In this evolutionary paradigm, bacteria are maintained in an energy-restricted environment in the growth phase called as long-term stationary phase or LTSP. This phase is characterized by a stable viable population size and highly dynamic genetic changes. Multiple independent iterations of LTSP evolution experiments have given rise to mutants that are slow-growing compared to the ancestor. Although the antagonistic regulation between rapid growth and stress response is fairly well-known in bacteria (especially *Escherichia coli*), the reason behind the growth deficit of many LTSP-adapted mutants has not been explored in detail. In this review, I revisit the trade-off between growth and stress response and delve into the regulatory mechanisms currently known to control growth under nutrient deficiency. Focusing on the theme of “sigma-factor competition” I try to search for the evolutionary reasoning of slow growth amongst mutants adapted to prolonged starvation. Additionally, I present novel experimental data indicating the dynamics of four such slow-growing variants that evolved during a 30-day long LTSP evolution experiment with *Escherichia coli*.

## 2. Introduction

Bacteria, one of the most resilient organisms on earth, are ubiquitous in our external and internal environments. In order to understand these organisms better, efforts to culture bacterial strains within the laboratory began in the early nineteenth century. In this regard, Louis Pasteur is credited with the formulation of liquid media, and Robert Koch with the invention of solid media (1). Over the next century, the metabolic and physical requirements needed for the rapid growth of a handful of microorganisms were standardized in the laboratory. However, in their natural habitats like soil, deep-sea surface, and within hosts, most bacteria have limited growth, mostly because of the lack of carbon, phosphate, and nitrogen sources in the ambient environment (2–4). The bacterial growth rate is an essential indicator of their adaptive fitness under ideal, non-stressful conditions, as demonstrated by decades-long evolutionary experiments (LTEE) with *Escherichia coli* (5–7). However, in the wild, acute scarcity of resources to build up biomass as well as competition from other organisms heavily limit bacterial growth. Hence, the opposing selection pressures to maximize growth and other survival modes (like eliciting general stress response pathways) dictate adaptive fitness in natural environments (2,3).

### 2.1 Slow-growth in mutants emerging from starved cultures: small-colony variants and their importance in public health

A large fraction of microbes occupying natural, nutrient-limited niches like soil, sections of the mammalian gut, and permafrost have been observed to be slow-growing (8–10). Slower growth in bacteria has been shown to be advantageous under carbon starvation (11). Within the confines of the laboratory, long-term evolution experiments conducted under prolonged starvation also led to the emergence of slow-growing mutants in both non-pathogens (12–14), and pathogens (15). This suggests a positive selection for slow-growth under long-term starvation. Very slow or near-dormant growth states have been demonstrated to increase fitness by enhancing tolerance to different stressors, e.g., antibiotic tolerance - a strategy exhibited by ‘persister’ populations in pathogens (16–18). Hence, slow-growth is an essential bet-hedging strategy deployed by a homogeneous population to survive dynamic and stressful environments.

Under extremely stressful conditions, some bacteria like *Bacillus spp* can form resilient structures called “spores” and remain in a non-growing, dormant state until favorable environment is restored, while other non-sporulating bacteria like *E. coli* exhibit very slow growth under stress (12,19). In *E.* c*oli,* growth rate is regulated by the antagonistic regulation balancing rapid growth and self-preservation (also called SPANC) (20). Whenever the ambient nutrient availability becomes sub-optimal, resources are allocated to express stress-response proteins as opposed to proteins assisting rapid growth (21), thereby reducing the growth rate.

A fraction of slow-growing bacteria are observed to form small-sized colonies on solid agar plates-a phenotype ubiquitously observed in specific isolates of both natural and laboratory-maintained strains. The first documented report of small colony variants(SCV) dates back to 1910 (22). Over time, more studies reported the emergence of small colonies upon exposure to chemicals like copper sulphate (23), phenol (24)), and antibiotics like gentamycin (18). SCV also poses a threat in the public health sector, as multiple groups demonstrated key pathogens to form SCV after being isolated from the site of infection, establishing SCV as a dominant phenotype among pathogens (25). *Staphylococcus aureus (26)* remains the most well-characterized species to form SCV, although other pathogens like *Pseudomonas aeruginosa* (18,27), *Staphylococcus epidermidis* (28), *Escherichia coli* (17), *Serratia marcescens* (29), and *Neisseria gonorrhoeae* (30) are also known to form small colonies. A reduced rate of respiration and pigmentation are characteristic of pathogenic SCVs (25). They consistently demonstrate slow growth primarily through metabolic mutations, either being auxotrophic to thymidine or being deficient in the electron transport chain (ETC) pathway. Due to their near-dormant metabolic state, SCVs may avoid immune responses from the host and multiple externally administered antibiotics (25) and persist within host niches for prolonged periods, driving chronic diseases (17). Enhanced tolerance of SCV to multiple stressors, a feature identified consistently even in non-pathogens, makes it a public health challenge. In order to specifically target and kill infectious SCVs further investigations into the regulatory origin of this phenotype are required both in the laboratory and clinical settings.

### 2.2 Repeated emergence of slow-growing variants during adaptive lab evolution in long-term stationary phase

Batch cultures of bacteria have been shown to remain viable for years without the addition of external nutrients in a paradigm called long-term stationary phase (LTSP) (31–34). In contrast to evolution under one or a set of pre-defined stressors, the evolving population is exposed to a dynamic set of stress factors during prolonged starvation. Under such conditions, batch cultures are shown to experience cycles of “feast-and-famine” in their environment akin to bacteria in natural habitats like the host, river bed sediment, or soil (12,32,35). Multiple independent studies have shown that the bacterial population increases in both genotypic and phenotypic diversity during evolution in batch cultures for prolonged periods (13,34,36). Slow-growing variants have also been observed to emerge during LTSP across different studies (12,13,34).

To check the repeatability of emergence of small colony variants in the LTSP paradigm, the LTSP evolution experiment was re-iterated following a previously published study from our group (36). As compared to the 5 strains that were evolved in the previous work (36), 11 evolving isolates were followed this time for 60 days (Fig 1). From these 11 isolates, SCV emerged in 4 lines, along with other interesting colony morphologies in the same sequence as compared to the previous runs (Fig 2). Across both runs of the evolution experiment, SCV was observed to emerge around the third week (Day 19-22 post-inoculation) of the experiment (Fig 1.A, Additional files 1), and was observed for two weeks until the cultures were monitored (Fig 1. B,C, Additional file 2). This suggests that underlying changes in media conditions select small colony phenotype during this period in prolonged stationary phase. The genetic basis of the SC described in Nandy *et al (13)* was mapped to the rpo operon, secifically - a point mutation in *rpoC* gene. Coding for the β’ subunit of RNA polymerase core, this gene is a fairly common target to be mutated across LTSP experiments (37). While analyzing the whole-genome sequences of the mutants from the second run of the evolution experiment, it was found that different SC isolates harbored unique mutations in different alleles and no alleles were featured in more than one strain (Table 1). No mutations were observed in the *rpo* operon as unique in any of the new SC strains, suggesting that slow growth is strongly selected in the LTSP but the genetic pathway to slow down growth is not fixed. In the second batch of SC mutants, mutations were also observed in several DNA-binding transcription factors like PaaX and Lon protease, some of which were previously implicated in survival under stress (38). Apart from global regulators, unique deletions in *acrB* a multi-drug efflux pump, and *cpdA*, the regulator of intracellular cAMP levels were found, indicating novel strategies to achieve slow growth (Table 1).

**Figure 1.**
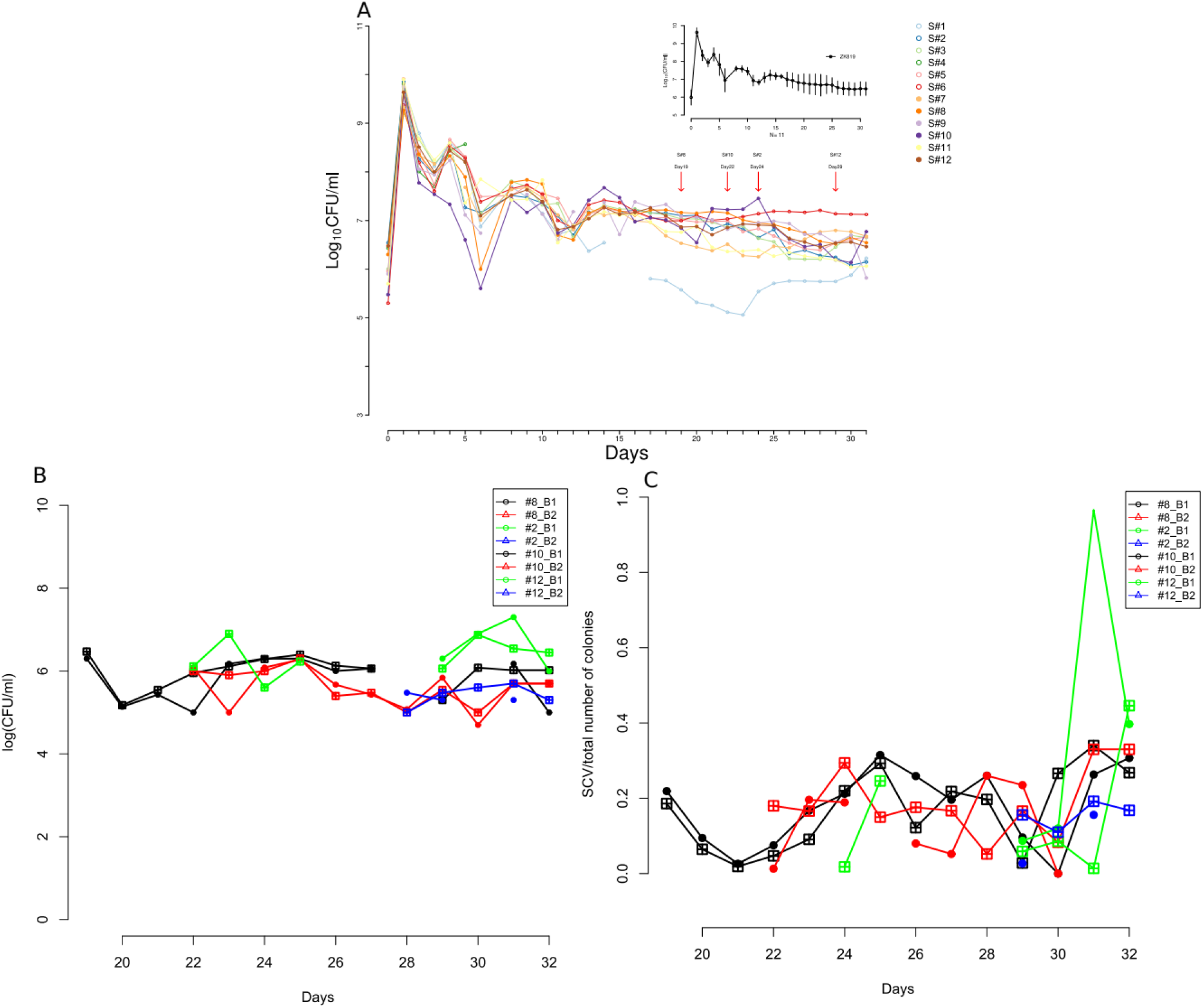
Evolution of replicate lines of Escherichia coli ZK819 in prolonged stationary phase. A. Colony count of 12 different replicates of E.coli ZK819 followed for 30 days in prolonged stationary phase in rich media. Inset: Average colony count across replicates over 30 days period. Standard deviation is represented as error bar at each data point. Red arrows indicate the days where small colony variants were first observed in different replicates B. Raw count of SC as observed in four independent replicates over last 12 days of evolution in prolonged stationary phase. C. Fraction of SC as observed on relevant days in 4 independent replicates (For raw data check Additional Files)

**Figure 2.**
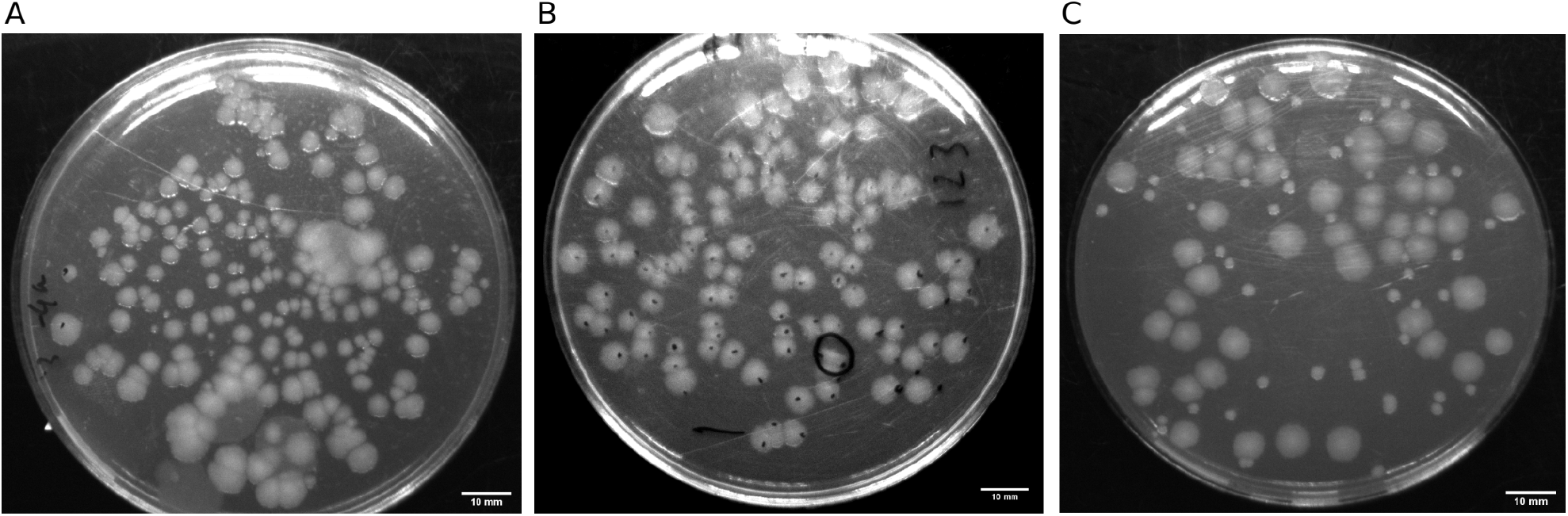
Representative images of different colony morphologies on LB plates after overnight growth during evolution prolonged stationary phase. A. Mucoid colonies shown by one independent line on Day 11. B. Fragmented colony(black circle) formed by one replicate on Day 15. C. Small sized colonies formed by one replicate line on Day 29

**Figure 3.**
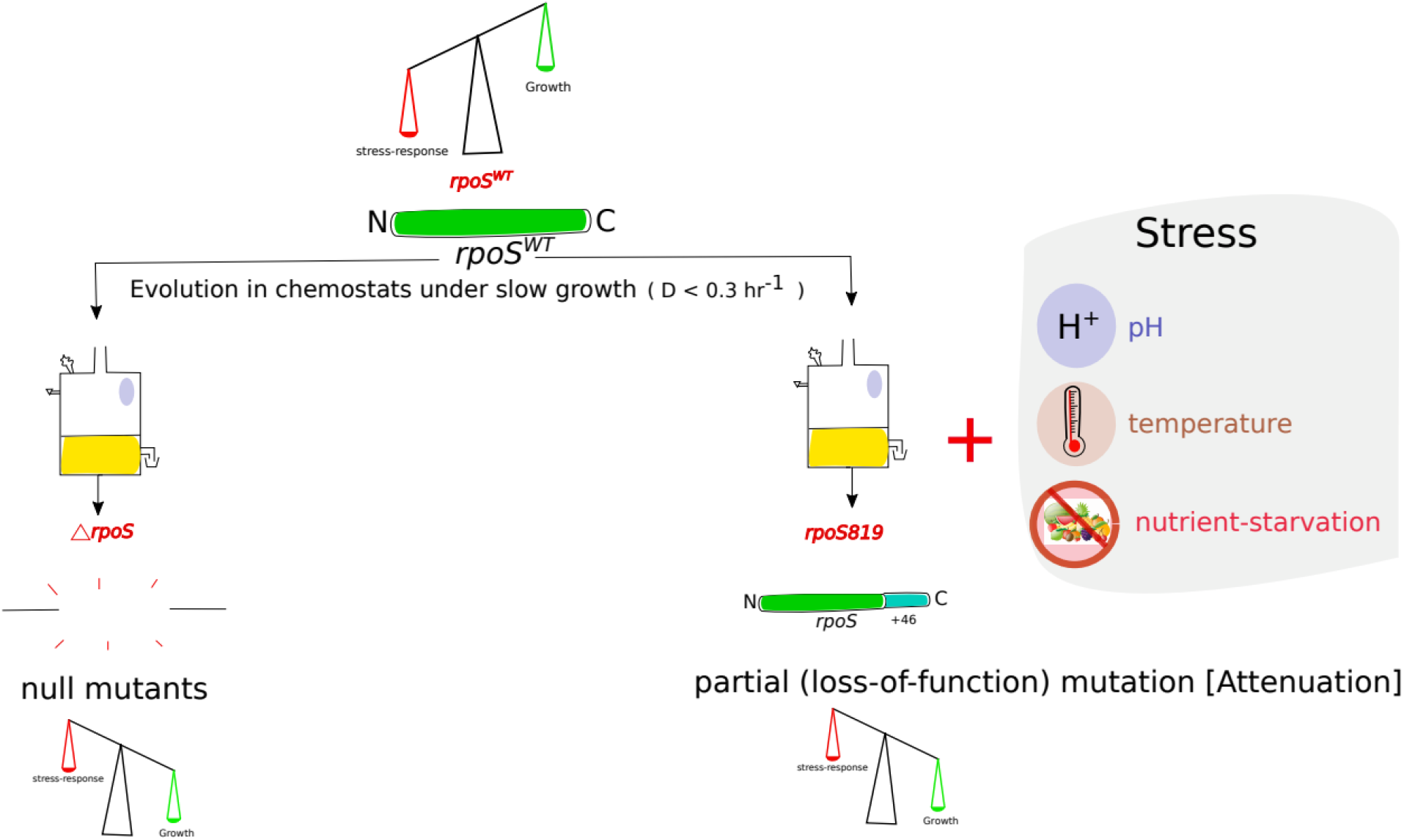
Change in the balance between growth and stress response modules in fast growing bacteria based on the evolution paradigm. Slow growth without the presence of additional stress factors results in complete loss of the stationary phase sigma factor allele rpoS. Presence of additional stress factors leads to the emegence of partial mutants of rpoS with lower efficiencies-that calibrates sigma factor competition.

**Table 1:** Unique mutation in SC strains of E. *coli* emerged from 1 month of starvation in LB.

| Strain of SC | Unique Mutation(s) |
| --- | --- |
| SC_sc | *rpoS*92, *rpoC*<sup>A494V</sup> |
| SC#2 | *gatA*(insertion of 1,199bp) |
| SC#8 | *paaX*(T59P), *lon*(L354Q) |
| SC#10 | *acrB*(Δ13 bp), *cpdA*(H164L) |
| SC#12 | *cpdA*(Q226*) |

Why are slow-growing variants so common across naturally occurring microbiota? Why are small colony variants repeatedly found to emerge from prolonged starved culture independent of ancestral genotypes? Although slow-growing strains are ubiquitously found in natural and lab-based strains under nutrient stress, this relation has not been summarily reviewed from a gene-regulatory and evolutionary perspective. Pursuance of this question deserves merit because most pathogens, as well as other ‘natural’ strains isolated from the wild, are generally slow-growing, and its evolutionary cause might hold insights into challenges like antimicrobial resistance, persistence - all of which are downstream effects of slow growth. In this review, I describe the existing knowledge of gene regulation under nutrient deficiency and focus on sigma-factor competition as a genetic mechanism that orchestrate bacterial growth under dynamic stress environments akin to those encountered in the long-term stationary phase.

## 3. Bacterial growth under prolonged nutrient limitation

Natural habitats are characterized by long periods of nutrient scarcity interjected by short stints of resource abundance termed as the “feast-and-famine” lifestyle (2,3,39). This metabolic cycle is thought to be the primary selection pressure in bacterial evolution (39). Within their natural niche bacteria spend most of their lifetime dividing with extremely low division rates under limited availability of carbon and nitrogen source, as opposed to extreme responses like the complete lack of growth (dormancy) or exponential growth (12,40). For instance, *E. coli* inhabiting the intestinal mucosa of the host are limited by carbon and nitrogen source and utilize complex sugars like N-acetyl glucosamide (NAG) and N-acetyl muraminic acid (NAM) to derive energy (25,41). Standard protocols for growing bacteria in rich media are unable to capture the genetic and transcriptional regulation that occurs under such nutrient-limited conditions (42). Multiple regulatory circuits have been evolved by bacteria to control its growth rate by constantly monitoring available resources under carbon, nitrogen-limited, and starvation conditions (43). In bacteria, rapid growth is facilitated by the housekeeping sigma-factor RpoD (*σ^D^)* whereas the starvation response comes under the ‘general stress response’ regulated by stationary-phase sigma-factor RpoS (*σ^S^)*. Tom Ferenci and colleagues have shown that hunger response and starvation response in bacteria are distinct and regulated based on the ambient nutrient levels (42). The *σ^S^*-controlled starvation response is triggered when the concentration of carbon source in the media falls below a concentration of 10-7 mM and is absent in both nutrient-excess (e.g, early and mid-exponential growth phases) and nutrient-limited (e.g, early stationary and prolonged stationary growth phases) conditions (20,40,42,44).

In this manner, the bacterial cell constantly monitors the nutrient status of its external environment and regulates its growth accordingly. In the following sections, I will discuss some of the mechanistic details that make the aforementioned regulation feasible.

### 3.1 *rpoS-*dependent strategies to control growth rate based on the ambient nutrientstatus

Initially, Richard Gourse and others reported different indications of gene regulation being controlled by the growth rate, primarily via regulation of the expression of *rrn* operons that control the intracellular ribosome levels (43,45). Global gene expression is modulated by the ambient growth rate by controlling the expression of genes involved in growth and cell division, and repressing the expression of other genes by sequestration of the core RNA polymerase via different sigma factors (43). Various sigma factors regulating rapid growth and general stress response compete to occupy the core RNA polymerase (46,47). As a result of this competition, different sets of genes are expressed based on which sigma factor is active, or bound to the core RNA polymerase. (43,48). Hence, binding of a particular sigma factor to the RNA polymerase core leads to upregulation of its own targets and downregulation of the target gene set of other competing sigma factor(s).

After bacterial cells exit the lag phase, gene expression is primarily modulated by *σ^D^,* promoting rapid growth because of the high ambient nutrient levels. As the population grows, the levels of carbon and nitrogen sources in the media steadily decline, compelling the bacterial cell to scavenge nutrients from the ambient media. This is achieved by activating the cellular “scavenging response” in two parts: (1) *changing the permeability of the outer membrane* through selective expression of porin channels *ompF* (higher permeability) or *ompC* (lower permeability) via the transcriptional regulator *ompR*, and (2) *overexpression of ABC type transporters* (*mgl, mal, LamB*) via an increase in the cAMP concentration (42). This “early stationary phase” response is prevalent at sugar concentrations ranging from 0.3 mM, when the growth rate is 70% of maximum growth rate, to 10^-6^ mM when growth becomes negligible (20,42). When the concentration of the carbon source drops below a threshold of 10-7mM, the growth-stress response balance could not be maintained by regulating only the membrane porosity, as most of the porins and transporter proteins are saturated in very low sugar concentration in the media. At this point, the *rpoS* or *σ^S^*-mediated “starvation response” is triggered.

The *σ^S^* mediated starvation response induces an array of changes like the compaction of the genome (49), reduction in the permeability of the cell membrane by expressing the low porosity porin *ompC* (50), increase in the production of osmoprotectants like trehalose by upregulation of *osmY* and *treA* (40), and repression of all genes directly promoting growth which helps bacteria to adapt to different stress factors that it endures in the stationary phase (46).

The ability of a bacterial cell to mount the *σ^S^*-mediated response is not devoid of cost, especially under prolonged periods of slow growth. This is evident from evolutionary dynamics in two different paradigms-a rapid decline of *rpoS^WT^* carrying variants in populations evolving in chemostats under low growth rates (D < 0.3 hr^-1^) (44,51), and selection of *rpoS* partial mutants in bacteria maintained in long-term stationary phase (52).

### 3.2 Balance between growth and stress response through sigma factor competition

Because of the dynamic nature of nutrient availability and presence of stressors in its environment, a constant balance has to be maintained between increasing biomass for rapid growth and self-preservation (46). Maintenance of this balance is crucial because over-optimizing rapid growth makes the population sensitive to even minute perturbations in the environment while putting excess weightage on self-preservation/stress-resistance results in the loss of fitness (46). It is known that *in vivo*, the concentration of the core RNA polymerase is limiting for transcription (46). Across different phases of bacterial growth, competition for the RNA polymerase core ensues between *σ^D^* and *σ^S^* to initiate transcription from specific promoters by integrating environmental cues (46). A change in the intracellular levels of any one of these sigma factors can indirectly affect the expression of genes under the control of the other one due to the competition as mentioned earlier (53). In E. *coli,* the intracellular levels of *σ^D^* have been shown to remain more-or-less constant across growth phases, whereas *σ^S^* concentration increases sharply during the onset of stationary phase (54).

It has been observed that strains carrying wild-type *rpoS* are able to utilize a lower number of carbon sources as compared to partial and null *rpoS* mutants *(20,55)*. Hence, *σ^S^* acts as a ‘necessary evil’ to the cell presenting a particular “cost” (lower flexibility in terms of carbon utilization) under nutrient-rich conditions, but becomes essential under stress (schematically shown in Fig. 3). It is not surprising that bacteria have evolved multiple regulatory circuits to control the expression and activity of *σ^S^* at all levels (comprehensively reviewed by Susan Gottesman and colleagues here: (56)).

### 3.2.1 Regulation of *rpoS* at different levels in the cell

At the sequence level, a wide variation is shown by *rpoS* across species (44,51,56,57). A variety of *σ^S^* partial mutants are selected from two separate evolutionary paradigms – 1) evolution in prolonged stationary phase, and 2) batch cultures maintained in chemostat under low growth rate and an additional ambient stressor (44,55,57). The ubiquitous observation of the emergence of *rpoS* mutants from several different evolution experiments underlines its utility in the diverse ecological niches (20). Transcription of the *rpoS* gene is controlled by multiple regulators, a key inducer being the small-molecule stress alarmone guanosine pentaphosphate or *ppGpp* (21,58,59). This regulatory role of *ppGpp* has broader implications for the competition between *σ^S^* and *σ^D^*, described in a later section of this review. Other known modulators of *rpoS* transcript levels are the small molecule regulator complex cAMP-CRP (60,61), and ArcA, the regulator of a two-component phosphotransfer system (62)

A stem-loop structure formed by the 5’-UTR region of the *rpoS* mRNA reduces its translation efficiency (63). A central role in the post-transcriptional modification and regulation of *σ^S^* is played by small RNAs like DsrA, RprA, and ArcZ, which bind to the 5’ UTR region. An adaptor protein called Hfq is necessary for these small RNAs to stabilize the nascent *rpoS* transcript (56). A fourth sRNA, OxyS, is known to negatively regulate *rpoS* levels, likely by occupying Hfq and out-competing other sRNAs (64). Many other genes like the *cspA* family (65), HU (66), and *csdA* (67) have been implicated in regulating *rpoS* mRNA stability by binding to the 5’ UTR.

Several elegant mechanisms have been evolved to control intracellular RpoS levels by regulating their rate of degradation. E*. coli* has a dedicated protease called ClpX to degrade the *σ^S^* protein (68). An adaptor protein called RssB is essential to identify RpoS and help in the function of the protease (69). RssB is identified to be the limiting factor for the degradation of RpoS, and its intracellular level is tighly controlled with the help of three anti-adaptor proteins - IraP, IraM, and IraD (70,71). These anti-adaptors are induced in response to environmental perturbations like DNA damage and cold shock primarily via the *ppGpp* mediated stringent response (56).

### 3.3. Heterogeneity in rpoS allele: nutrient scavenging as an active strategy to survive nutrient limitation

Initial studies characterizing long-term cultures mostly relied on metabolic readout via growth rates and phenotypic annotation of various facets of the bacterial lifestyle. In recent years the real-time monitoring of the mutation frequencies over prolonged periods has been enabled by the advent of massively parallel high-throughput sequencing. Identification of multiple subpopulations with unique genomic signatures is now possible by combining adaptive lab evolution with periodic population sequencing of evolving isolates (5,34,36,72,73). Across these experiments, *rpoS* has emerged as a prime target for mutation, allowing bacteria to adapt to different stress factors in their environment.

#### 3.3.1 *rpoS* mutations that emerge under slow growth in chemostats

Chemostats allow the continuous monitoring of cultures growing with a specific growth rate (exponentially) by regulating the efflux of nutrients via the dilution rate (D). Mutations in multiple global regulators like *rpoS* are found in strains evolved in chemostats under glucose limitation (74). In another study, mutations in *rpoS* were quickly accumulated by E. *coli* growing with growth rates ≤0.3 hr^-1^, and the proportion of cells carrying wild-type *rpoS* rapidly declined in carbon and nitrogen-limited media (44). Both partial and null mutants of *rpoS* emerged with a specific pattern based on the period of stress exposure, with slow growth under carbon and nitrogen limitation largely giving rise to loss-of-function mutants of *rpoS*, while nutrient limitation coupled with other stress factors like pH selecting for partially attenuated *rpoS* mutants *(44,46)* (Fig. 3). This highlights the role of ambient nutrient status directly shaping the bacterial genomes by selecting mutations in global regulators which in turn regulate a large number of smaller regulatory circuits or local regulators.

### 3.4 Regulatory programs to modulate sigma-factor competition

Multiple independent regulatory circuits have been demonstrated to influence the competition between *σ^D^* and *σ^S^*, thus regulating the balance between self-preservation and nutritional competence (SPANC) in response to the ambient nutrient and stress status (20,56). Some of these regulators increase the efficiency of *σ^S^* to bind to the RNA polymerase core, while others interfere with the binding of *σ^D^* with the core polymerase and hence repress its activity. In this review, I will focus on the role of a few global regulators essential for survival under prolonged nutrient limitation.

### 3.4.1 *ppGpp:* the master regulator of sigma factor competition

One of the most important reporters of ambient carbon and nitrogen levels that controls global gene expression in bacteria is the small molecule stress alarmone guanosine pentaphosphate or *(p)ppGpp* (53). The intracellular levels of this modified nucleotide govern the rapid growth versus self-preservation balance. During exponential growth, the concentration of *ppGpp* in the cell remains low due to the nutritional abundance. During starvation periods, free/stalled ribosomes resulting from a lack of amino-acyl t-RNAs induce the production of *ppGpp* via the expression of *relA* and *spoT* (53). *ppGpp* functions by destabilizing the DNA-protein interactions leading to the dissociation of *σ^D^* from the RNA polymerase holoenzyme. This event increases the availability of core polymerase for *σ^S^* to bind. *σS-*bound holoenzyme can then transcribe from cognate promoters that encode genes involved in stress response (59). In fact, the expression and activity of *σ^S^* are themselves regulated by *ppGpp,* establishing the latter as the “master regulator” of sigma-factor competition (46,53,75).

### 3.4.2. Role of *Crl* and *rssB* in governing sigma factor competitions

Crl is a global regulator, active during the transition from exponential growth to stationary phase, that positively affects the activity of *σ^S^* by facilitating the binding of RpoS with the core RNA polymerase (76)(56,77). In effect, the sigma factor competition between *σ^S^* and *σ^D^* for the core RNA polymerase is shifted by Crl in favor of *σ^S^* - firstly by increasing the expression of a large subset of the *σ^S^* regulon under low RpoS concentrations, and secondly by inducing proteolysis of RpoS via increased expression of RssB (69,77). Hence regulation by Crl leads to a lesser but more active *σ^S^* protein which indicates that some *σ^S^* targets are probably expressed during exponential growth (77). This regulation has implications in the log-to-stationary transit in batch cultures and during the long-term evolution of starved cultures (77).

6S RNA, a small, non-coding RNA generated from *ssrB* gene is also instrumental in controlling sigma factor activity (56,78). This RNA binds to the *σ^D^*-core polymerase complex by mimicking a promoter and inhibits transcription from several *σ^D^* promoters, and also allows *σ^S^* to occupy the free core polymerase and transcribe from relevant targets (78,79). The 6S RNA is shown to regulate different sets of genes across various growth phases, indicating that its activity is governed by the ambient nutrient status (79). Another protein Rsd upregulates a subset of *σ^S^* regulon by directly sequestering free *σ^D^* and binding to the core polymerase enzyme, effectively shifting the sigma factor competition in favor of *σ^S^* (80). The cross-talk of these individual regulators enables the cell to fine-tune the SPANC balance by monitoring ambient nutrient and stress states. Rsd and 6S RNA are found to repress each other via transcriptomic analyses in *ΔssrS* and *Δrsd* knockouts (79). Further 6S RNA also controls the expression of Crl, and regulates the sigma factor competition likely by altering the levels of free and polymerase-bound *σ^D^* in the cell (79).

Finally, the sigma-factor competition is affected by the general nutrient status of the media and the stress factors present in the ambient environment. This competition regulates the pattern of global gene expression in the bacterial cell, and hence multiple mutations selected under nutrient stress fine-tune these dynamics to orchestrate the balance between rapid growth and stress response (46).

### 3.5 Growth Advantage in Stationary Phase (GASP): conferring fitness by modulating sigma factor competition

The niches naturally occupied by bacteria in terms of the nutrient status and ambient stress factors are closely mimicked by the long-term stationary phase (LTSP) (37). During growth in LTSP, younger cultures are outcompeted by aged cultures via a phenotype termed as “growth advantage in stationary phase” or GASP. This phenotype is heavily modulated by environmental factors like the genetic background of the competing strains, and the nutrient status of the media (13,81,82). GASP is also controlled by physical factors like the volume of the culture, shaking frequency of the culture, and even the shape of the conical flask carrying the culture (83). Investigations by different groups on the genetic basis underlying GASP revealed the involvement of the global regulators like different sigma factors, the amino acid metabolism regulator *lrp* (84), specific protein carboxyl methyltransferase (85), the iron-binding protein Dpr (86), and the structural subunits of RNA polymerase, coded by the *rpoABC* operon (13,36,87) in the process. However, the first identified and most studied GASP mutation remains an allele with a 46 base-pair duplication at the C-terminal end of the *rpoS* gene, resulting in an elongated and attenuated sigma factor termed *rpoS819* (33,47). Recently, it was shown that in the initial stages of LTSP evolution, mutations in global regulators confer a higher fitness advantage- an effect that declines over time, leading to local regulators being mutated at a higher frequency later in the experiment (88). Evidence and role of GASP have been detected in pathogen populations as well-vector-borne pathogens like X. *nematophila* balance a trade-off between pathogenicity and the GASP phenotype, with the competitively advantageous *lrp* mutant subpopulations lacking transmissivity and virulence later in evolution (89).

#### 3.5.1 Genetic determinants of GASP: mutations in the stationary phase sigma factor(*σ^S^*) and RNA polymerase complex

Heterogeneity of *rpoS* allele across bacterial species and its variation in the expression levels have been described across multiple studies (44,51,57,90). Recently, it was shown that *rpoS819* suffers a re-duplication of the 46 base-pair region when passaged in LTSP, forming an even more elongated allele *rpoS92* which can confer GASP and has efficiency closer to the *rpoS^WT^* allele (13,60).

The basal expression of *rpoS* is optimized based on nutrient availability and stress factors in the ambient niche and is observed to differ between different strains of the same species. This variation contributes to phenotypic heterogeneity-possibly as a bet-hedging strategy to survive unfavorable conditions (20,55). Functional attenuation of *rpoS* leads to the super-induction of *rpoD* regulon, resulting in the de-repression of specific rpoS-induced high-affinity transporters, increasing fitness advantage under carbon limitation (42). *rpoS* null mutants are selected under only nutrient stress (D < 0.3 hr-1), but when additional stressors (like pH, temperature) are present, partial mutants (with lower expression/efficiency) are selected (44).

Apart from *rpoS*, LTSP evolution experiments starting with different ancestors and media by different groups have also identified mutations in various RNA polymerase core subunits (ααββ’γω) and global regulators like *crp*, *cpdA*, and *fusA* (36,87,91). Although the precise mechanism by which these different mutations increase survival in long-term starved cultures is not completely understood yet, analysis of converging mutations across different ancestor strains reveals selection pressures that are active in LTSP. An evolution experiment maintaining E. *coli* MG1655 strain in the prolonged stationary phase was carried out by the Hirschberg group for three years and the dynamics of different mutations in the first four months (91) and cumulatively over three years (34) was reported. 90% of all clones sampled over six different timepoints in their experiment carried a substitution in *rpo* operon coding for the structural subunits of RNA polymerase, and most of these mutations were in one of three loci, viz, *rpoB* 1272, *rpoC* 334, and *rpoC* 428 (91). These mutations emerged within the first four months of the evolution experiment and persisted till three years into starvation (34). In another LTSP experiment with E. *coli* K-12 *str* ZK819 as the ancestor, *rpoABC* mutations were found to be widespread among the variants that were sequenced at different timepoints for a month(36). Most of these alleles are found to decrease growth rate in the exponential phase and hence present a cost to the bearing cell (13,91). However, the selection of these mutations across different ancestors and runs of LTSP evolution point to the obvious advantage of slow growth in nutrient-constricted spent media. Indeed, media age has emerged as a major determinant of the competitive fitness of LTSP-adapted mutants (13). Likely, the balance between rapid growth and stress response (SPANC) will be affected by the growth defect acquired by mutations in the *rpoABC* operon. Diverse strategies to gain fitness under prolonged starvation are developed by the bacterial cell by fine-tuning this SPANC balance (55)(Fig. 3).

Apart from the *rpo* operon, other genes that are frequently mutated in LTSP adapted variants include the cyclic-AMP associated global regulated CRP, cAMP phosphodiesterase *cpdA,* and the translational elongation factor *fusA* (EF-G). All of the above-mentioned genes hold the potential to modulate the SPANC balance via controlling the sigma factor competition.

Apart from the above-mentioned factors, mutations in other sigma factors like *σA* in *Bacillus subtilis* and *σN* in E. *coli* are also reported to alter the SPANC balance, enabling the bacterial cell to respond to environmental stressors (92,93). Victoria Shingler has proposed a theory where the binding between *σ^D^* and RNA polymerase core is affected by mutations in *σ^D^*, or concomitant players like alternative sigma factors and core RNA polymerase subunits. This increases the availability of free core polymerase to be occupied by other ‘non-housekeeping’ sigma factors, likely via the involvement of small-molecule regulators like *ppGpp* (75). Evidence of this theory has been provided by work done in Ruth Hershberg and Seshasayee groups.

## 4. Conclusion

### 4.1 The path ahead: Challenges in studying the long-term stationary phase

The long-term evolution of bacteria in chemostats (under constant nutritional status) has been studied for more than 20 years (94). The short generation time of a few species has enabled these seminal studies to uncover general principles about bacterial gene regulation under different growth rates and how it shapes the genome over time (95,96). However, a constant supply of external nutrients at a pre-determined rate is required in this paradigm. In contrast, evolving strains do not require constant input of nutrients in the ‘long-term stationary phase’ paradigm. This entails minimal perturbation to the system by the observer, leading to LTSP being acknowledged to be closest to the environment encountered by bacteria in the wild (32,47). The earliest evidence of GASP was reported by Roberto Kolter’s group in 1993 that identified a novel allele of *σ^S^*, termed *rpoS819*, to be responsible for competitive fitness (33). Following this study, further key GASP mutations that emerge contingently under the *rpoS819* background were uncovered, the most important of them being the global regulator *lrp* (leucine receptor protein) and the *ybej-gltJKL* cluster (84,97,98). Although the gene regulatory mechanisms leading to GASP have been identified and fairly well-studied, the exact metabolic basis for the same is not yet known. Most of the known causative mutations conferring GASP are mapped in global regulators and cause large-scale alterations across the transcriptome. This makes it difficult to comment on the metabolic capacities of any sequenced GASP mutant. As previous studies have shown that the GASP phenotype is heavily dependent on the nutritional status of the ambient media (13), chemical characterization of differently aged spent/nutrient-depleted media is essential. However, this has proven difficult to achieve via untargeted mass spectrometric methods because of the inherent variability across rich media. Novel approaches need to be developed to characterize a wide variety of novel metabolites from rich media, as novel amino acids have already been isolated from spent media (98).

Multiple other looming questions are yet to be answered in this field-what are the mutation landscapes across genomes that enable GASP? How is the emergence of GASP-enabling mutations dependent on genetic contingency and environmental parameters? How are real-world situations like host-pathogen interactions affected by GASP and LTSP-induced stress tolerance?

Finally, LTSP has been demonstrated as a useful paradigm to discover bacterial strategies to survive in a severely nutrient-depleted environment, akin to what they face within the host system during infection and in diverse natural niches. Recently the competition between two E. *coli* sigma factors controlling growth and stress response were identified as a strategy to gain a transient competitive advantage in LTSP (13). In LTSP, the genotypic and phenotypic variation of the initial, genetically homogeneous culture increase, generating subpopulations that compete amongst themselves to survive and thrive (32,36) by recycling the debris left by the previous subpopulations (99). Similarly, persistor subpopulations that can endure antibiotic regimes and drive chronic diseases are formed by pathogens in nutrient-limited host niches (10,100). Resistance to novel non-exposed stress factors in evolved mutants has been identified in LTSP evolution experiments (13,101). The reason for gaining these novel resistances is not clearly understood currently, but it might be caused by a global up-regulation of the general stress response (GSR) pathway. The translational utility of the LTSP paradigm can be harnessed by evolving pathogens in conditions resembling specific host niches and identifying strategies developed by the strain to gain fitness (89,102).

Bacterial gene regulation in long-term stationary phase is dynamic, complex, and spread across multiple subpopulations in an ever-changing environment. The role of external factors in the evolutionary dynamics in this phase is being increasingly recognized. Although the idea of sustained cultivation of microbes is not new, systematic observation of genomic and transcriptomic changes in this survival phase has only been started in the last 30 years (Fig 4). LTSP being a temporally open-ended paradigm, the amount of information that could be mined via novel, high-throughput multi-omics tools is endless. The natural evolutionary dynamics in biomes like host gut and soi are closely mimicked by the LTSP paradigm since it is free from any perturbation by the experimenter. Further developments against current medical challenges like the evolution of multi-drug resistance (18), de-novo stress resistance (101), and phenotypic persistence (100) could be made by exploring the eco-evolutionary dynamics in LTSP.

**Figure 4.**
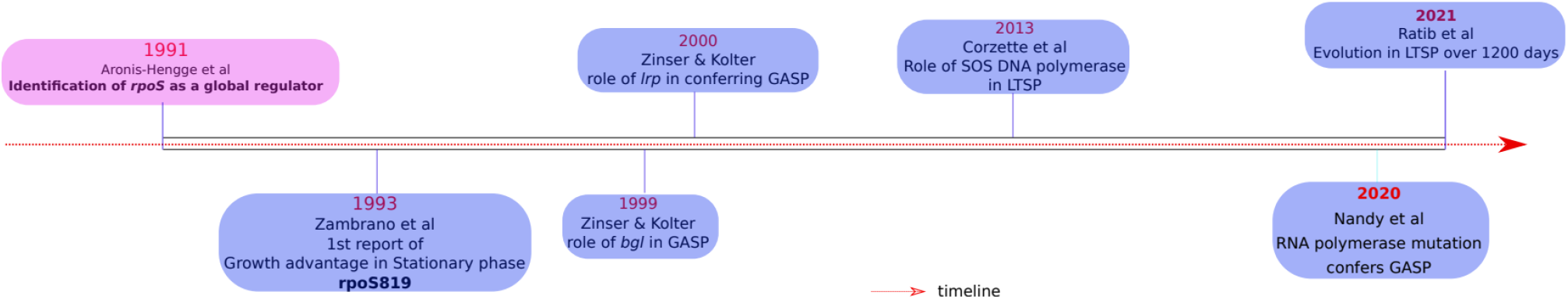
A timeline showing the key developments in the field of LTSP evolution and bacterial gene regulation

## 5. Declarations

### 5.1 Ethics approval and consent to participate

Not applicabe for this work

### 5.2 Consent for publication

Not applicable for this work

### 5.3 Availability of data and materials

The whole genome data analysed for the mutations in Table 1 is already uploaded in NCBI Sequence Read Archive (SRA) under the accession number SRP236016 (https://trace.ncbi.nlm.nih.gov/Traces/sra/?study=SRP236016). The colony count data acquired from the long-term stationary phase evolution experiment are available in this manuscript as Additional Files.

### 5.4 Authors and Contributors

This manuscript was conceived, drafted, and written by PN. All the raw data described in the manuscript was acquired and analysed by PN.

## 5.7 Acknowledgements

The author wants to thank Dr. Aswin Sai Narain Seshasayee, Dr. Deepa Agashe, and Dr. Laasya Samhita for providing critical feedback and improvising the manuscript.

## Additional/Supplementary file information

- <u>File name = Additional File 1</u>
- File type = Comma-separated value (csv) file
- Title = Daily colony count data for 12 replicates of Escherichia coli evolved in long term stationary phase
- Description = Raw colony count, taken at a daily frequency from 12 biological replicates of *Escherichia coli* ZK819, maintained under prolonged starvation in LB for 30 days. Further, a separate count of small colonies are kept as and when observed across four different lines

- <u>File name = Additional File 2</u>
- File type = Comma-separated value (csv) file
- Title = Fraction of Small colony variants across different evolving lines of *Escherichia coli* ZK819 under prolonged starvation
- Description = This file lists the relative frequency of small colony acorss two technical replicate platings from each evolving lines of *Escherichia coli* ZK819 maintained under long term starvation. Average counts are derived from two replicate readings everyday where SCV could be observed, and the error is expressed as standard deviation of the two readings.

